# Single-sample, multi-omic mass spectrometry for investigating mechanisms of drug toxicity

**DOI:** 10.1101/2025.02.13.638125

**Authors:** Iqbal Mahmud, Wai-Kin Chan, Karen Yannell, Cate Simmermaker, Genevieve Van de Bittner, Linfeng Wu, Daniel Chan, Sheher Banu Mohsin, Yiwei Liu, John Sausen, John N. Weinstein, Philip L. Lorenzi

## Abstract

Poor therapeutic index is a principal cause of drug attrition during development. A case in point is L-asparaginase (ASNase), an enzyme-drug approved for treatment of pediatric acute lymphoblastic leukemia (ALL) but too toxic for adults. To elucidate potentially targetable mechanisms for mitigation of ASNase toxicity, we performed multi-omic profiling of the response to sub-toxic and toxic doses of ASNase in mice. We collected whole blood samples longitudinally, processed them to plasma, and extracted metabolites, lipids, and proteins from a single 20-µL plasma sample. We analyzed the extracts using multiple reaction monitoring (MRM) of 500+ water soluble metabolites, 750+ lipids, and 375 peptides on a triple quadrupole LC-MS/MS platform. Metabolites, lipids, and peptides that were modulated in a dose-dependent manner appeared to converge on antioxidation, inflammation, autophagy, and cell death pathways, prompting the hypothesis that inhibiting those pathways might decrease ASNase toxicity while preserving anticancer activity. Overall, we provide here a streamlined, three-in-one LC-MS/MS workflow for targeted metabolomics, lipidomics, and proteomics and demonstrate its ability to generate new insights into mechanisms of drug toxicity.

## INTRODUCTION

Elucidating mechanisms of drug toxicity remains a critical challenge in drug development^1^. Here, we sought to develop a multi-omic, system biology approach using mass spectrometry (MS)-based profiling, with the goal of generating new insights into the complex molecular networks that underlie drug-induced adverse effects^2^. However, technical limitations in sample preparation and multi-omic data integration have historically constrained such approaches^3^. Here, we present a streamlined, automated sample preparation and MS-based workflow that enables simultaneous characterization of metabolites, lipids, and proteins from minimal sample volumes.

We demonstrate the method’s utility in the context of L-asparaginase (ASNase)—an enzyme-drug used to treat pediatric acute lymphoblastic leukemia (ALL)^4^. It catalyzes the enzymatic degradation of asparagine, targeting cancer cells that exhibit asparagine dependency^5^. Despite over half a century of research on ASNase, major challenges remain^6^. For example, the *in vivo* pharmacodynamics (PD) of ASNase could not be assessed accurately until we and others developed methods to overcome significant analytical challenges^7^. Additionally, the contribution of the glutaminase activity of ASNase to its overall anticancer activity was unclear, and we demonstrated that the glutaminase activity is required for anticancer activity^8^. A current challenge is that ASNase-induced toxicities, including hyperammonemia^9^, hypersensitivity^10^, pancreatitis^11^, thrombosis^12^, hepatotoxicity^13^, hyperglycemia^14^, hyperbilirubinemia^14^, and hypertriglyceridaemia^15^ limit the use of ASNase in adults.

Here we report a multi-omic LC-MS/MS workflow to elucidate mechanisms and biomarkers that could potentially be targeted to mitigate ASNase toxicity. The resulting workflow enables: 1) measurement of metabolites, lipids, and proteins from a single 20-µL sample of plasma, facilitating longitudinal multiomic studies in mice; 2) association of candidate biomarkers with drug toxicity and efficacy; and 3) identification of potential combination therapy targets. While we developed and validated this drug toxicity workflow specifically for mouse plasma samples, the underlying analytical principles and sample preparation strategies suggest potential utility for multi-omic analyses in other experimental systems, though additional validation would be required.

## EXPERIMENTAL SECTION

### Sample collection

The research complies with all relevant ethical regulations: MD Anderson Cancer Center IACUC-approved protocol 00001658-RN02. NSG mice [NOD.Cg-PRKDC(scid) IL2RG(tm1Wjl) (The Jackson Laboratory stock #005557) housed in a pathogen-free vivarium were treated with either PBS (vehicle, n = 3), Spectrila® (recombinant *E. coli* ASNase) at 2,500 IU/kg (n = 4), Spectrila® at 10,000 IU/kg (n = 3), or Spectrila® at 20,000 IU/kg (n = 3). Based on our previous work, the two highest doses are known to be toxic, and 2,500 IU/kg is known to be sub-toxic. Treatments were administered intraperitoneally at a volume of 100 µL every day for 14 d.

Approximately 150 µL whole blood was collected from the submandibular vein into a K2-EDTA micro capillary blood collection tube (RAM Scientific #07-7052) on day -1 (one day before treatment) and posttreatment days 7, 15, and 20. Blood was centrifuged at 2,000 *g* for 15 min at 4°C. Plasma supernatant was promptly transferred into a clean polypropylene tube, aliquoted, and stored at -80°C.

### Multi-omic (3-in-1) sample preparation

The Captiva EMR-Lipid sample filtration plate provides highly selective, efficient lipid/removal based on size exclusion and hydrophobic interaction. Enhanced matrix removal (EMR) minimizes ion suppression of target analytes, significantly improving method ruggedness and precision. The 96-well plate and 1 mL cartridge formats contain a solvent retention frit, which allows in-well protein precipitation that streamlines sample preparation. An AssayMap Bravo Sample Prep Platform was used to automate separation of metabolites, lipids, and proteins from 20 µL plasma. Protein samples further underwent trypsin digestion and desalting on the Bravo. Metabolite and lipid fractions were vacuum concentrated to dryness and reconstituted in the mobile phase A corresponding to each workflow..

### LC-MS/MS for Quantitative Multi-Omics

Liquid chromatography-tandem mass spectrometry (LC-MS/MS) methods were developed on the Agilent 6495D triple quadrupole with 4^th^ generation iFunnel technology (combination of highand lowpressure funnel). The method achieves high reproducibility at ultra-low dwell times, enabling highplex multi-omics in a targeted fashion. A 1290 Infinity II Bio LC front end is lined with a bio-inert material, MP35N, that decreases metal ion interactions with metal-sensitive analytes, providing lower background, better peak shape, and better sensitivity.

### Proteomics

MRM Proteomics kits containing stable isotope-labeled peptide internal standards were leveraged for the targeted measurement of 375 proteins. A ZORBAX RRHD Eclipse Plus C18 column, 2.1 × 150 mm, 1.8 µm particle size (p/n 959759-902) was used for separation. Mobile phase A contained H_2_O with 0.1% formic acid (FA), and mobile phase B contained acetonitrile with 0.1% FA. Flow rate was 0.4 mL/min, and injection volume was 10 µL. The gradient was provided with the MRM Proteomics kit: 0 min 2%B, 2 min 7%B, 50 min 30%B, 53 min 45%B, 53.5 min 80%B, 55.5 min 80%B, 56 min 2%B, 60 min 2%B.

### Metabolomics

The metabolomics workflow used a Poroshell 120 HILIC-Z column (2.1 × 150 mm, 2.7 µm particle size, Agilent # 683775-924) and column temperature 15°C. Mobile phase A was 20 mM ammonium acetate, pH 9.3 with 5 µM medronic acid in water, and mobile phase B was pure ACN. Flow rate was 0.4 mL/min, and injection volume was 1 µL. The gradient was 0 min 90% B, 1 min 90%B, 8 min 78%B, 12 min 60%B, 15 min 10%B, 18 min 10%B, 19 min 90%B at 0.5 mL/ min, 23 min 90%B. A metabolite MRM database containing over 500 annotated metabolites was leveraged to find 314 metabolites in this plasma study.

### Lipidomics

For lipidomics we used a Zorbax Eclipse Plus C18 column (100 × 2.1 mm, 1.8 µm particle size, Agilent #959758-902) and column temperature 45°C. Mobile phase A contained 50% water, 30% acetonitrile, 20% isopropanol, 10 mM ammonium formate, and 5 µM medronic acid. Mobile phase B contained 1% water, 9% acetonitrile, 90% isopropanol, and 10 mM ammonium formate. Flow rate was 0.4 mL/min and injection volume was 1 µL. The gradient was 0 min 15%B, 2.5 min 50%B, 2.6 min 57%B, 9 min 70%B, 9.1 min 93%B, 11 min 96%B, 11.1 min 100%B, 12 min 100%B, 12.2 min 15%B, 16 min 15%B. Lipids were identified using a curated lipid MRM database featuring extensive double bond annotation containing over 750 lipids.

### Data analysis and interpretation

Raw data was imported into MassHunter Quantitative Analysis (v 12). MRMs were extracted and integrated by Agile2 integrator. Area counts were exported into excel for import into statistical workflows. Multivariate analysis and metabolite set enrichment analysis was performed using MetaboAnalyst 6.0^16^. Newt 4.0^17^ System Biology Graphical Notation (SBGN)-based biological maps were built for multi-omic data enrichment analysis.

## RESULTS AND DISCUSSION

### Streamlined multi-omic analysis exhibits rigor and reproducibility

The optimized workflow involves the sequential extraction of metabolites, lipids, and proteins from a single 20 µL plasma sample (**Figure 1A**). Within the 314 metabolites, 750 lipids, and 375 proteins detected in our samples, 81%, 99%, and 91%, respectively exhibited less than or equal to 20% RSD (**Figures 1C-E**). The combined depth of coverage and quantitative rigor achieved by the workflow blur the line between multi-omic discovery and validation, enabling validation-grade quantitative rigor in a discovery workflow.

**Figure 1.**
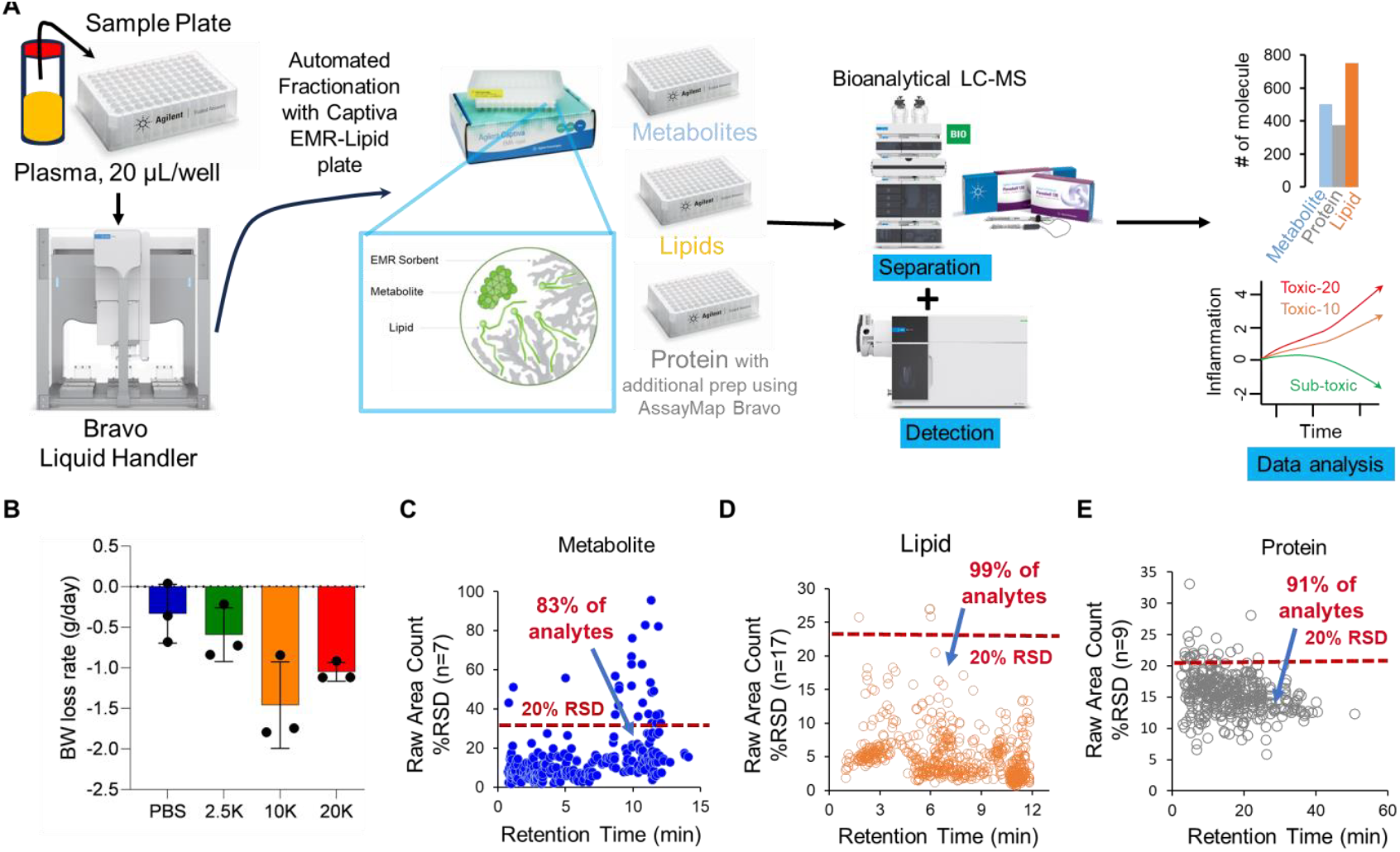
Bioanalytical LC/MS-based multi-omics workflow. **(A)** The workflow starts with a single plasma sample and serial extractions to separate components, including metabolites, lipids, and proteins. Body weight loss is used as an index of ASNase toxicity (**B**). Compound reproducibility across the retention time (RT) range for metabolite (B), lipid (C), and protein (D), respectively.

### Multi-omic analysis facilitates discovery of biomarkers associated with ASNase toxicity

Next, we analyzed metabolomic, lipidomic, and proteomic data obtained from mice treated with increasing concentrations of ASNase.

Multivariate receiver operating characteristic (ROC) curve analysis yielded AUCs of 0.869, 0.917, and 0.914 for metabolites, lipids, and proteins, respectively **(Figures 2A-C**). Integration of all three data types yielded an AUC of 0.95 (**Figure 2D**), indicating that multi-omic data improved the sensitivity and specificity for detection of ASNase toxicity compared to any of the individual omic data types. Principal component analysis (PCA) confirmed distinct metabolomic, lipidomic, and proteomic profiles associated with sub-toxic and toxic doses of ASNase (**Figures 3A-C** and **Figures S1-S4**).

**Figure 2.**
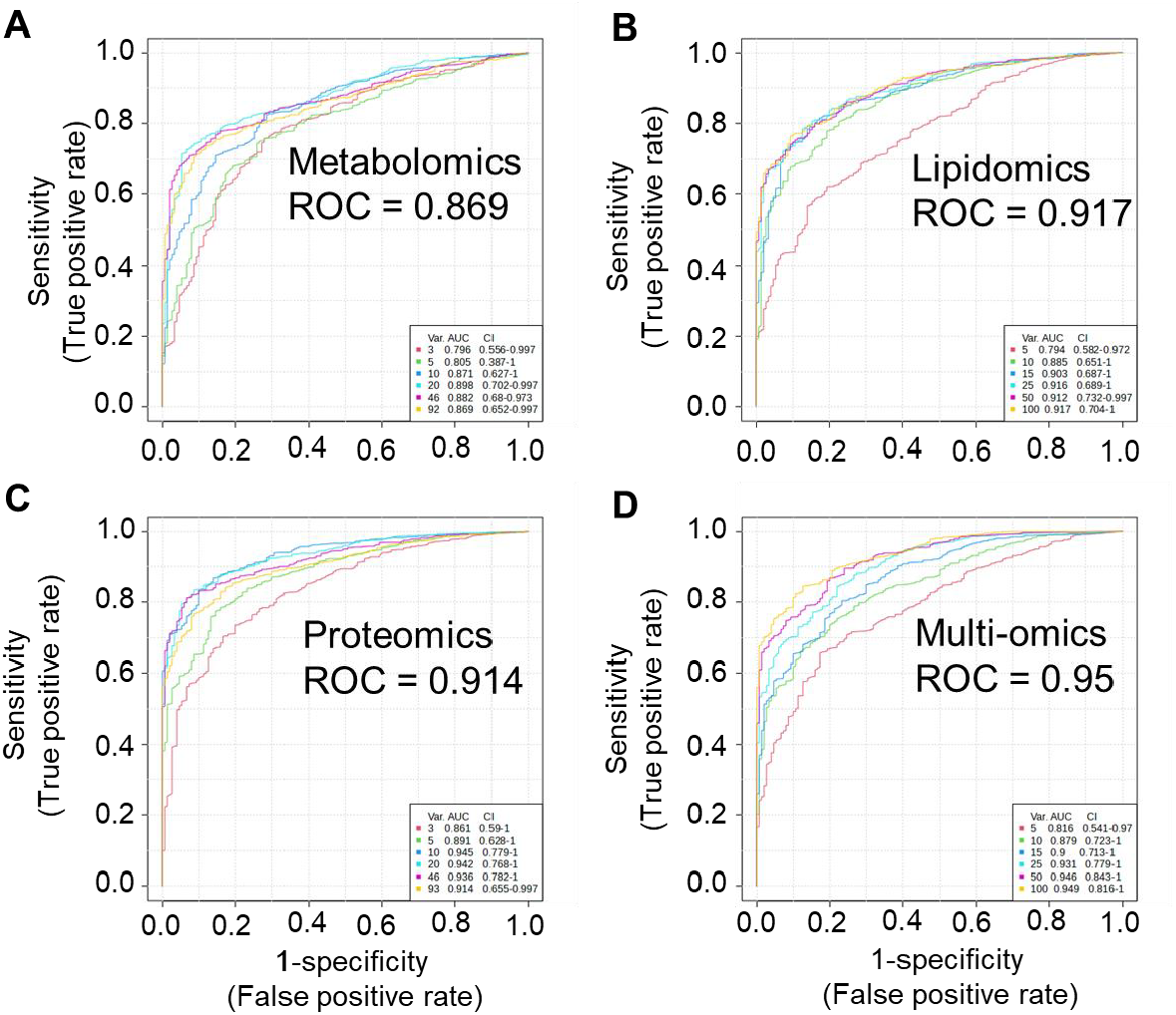
Biomarker prediction by multivariate ROC analysis. Shown are the area under the curve (AUC) analyses of n = 9 untreated and n = 30 drug-treated cell samples for metabolomics (A), lipidomics (B), proteomics (C), and multi-omics (D), respectively.

**Figure 3.**
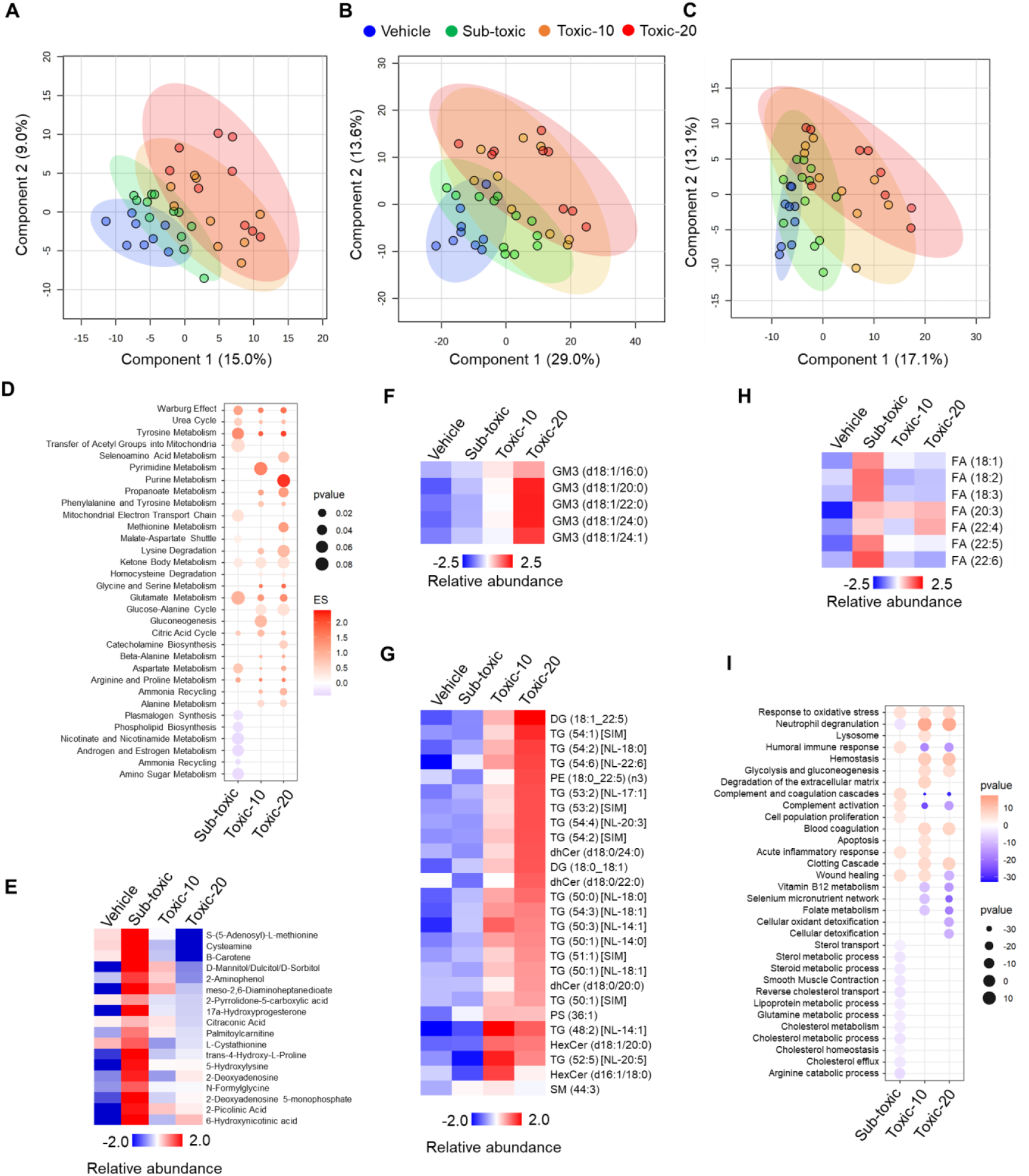
Dose-dependent modulation of metabolites, lipids, and proteins by ASNase reveals potentially targetable pathways for mitigating ASNase toxicity. Multivariate cluster analysis of metabolomic (**A**), lipidomic (**B**), and proteomic (**C**) profiles for vehicle and different doses of ASNase. Significantly modulated pathways are shown for metabolite (**D, E**), lipid (**E**,**F, H**), and protein (I, **G**), respectively.

We next compared ASNase-treated and PBS-treated metabolomic, lipidomic, and proteomic results to investigate mechanisms of toxicity. Analytes with an average toxic:sub-toxic ratio greater than or equal to 1.5 and a p-value < 0.05 were classified as differentially modulated. The sub-toxic dose (2,500 IU/kg) yielded down-regulation of metabolites associated with hormone and nitrogen metabolism (**Figure 3D**). The sub-toxic dose also up-regulated glutamate and glutathione networks and metabolites associated with mitochondrial function and antioxidation. Those are likely adaptive responses to counteract ASNase effects (**Figure 3D**). We observed inverse trends following treatment with toxic doses (10,000 and 20,000 IU/kg) of ASNase. The observed downregulation of S-(5-adenosyl)-L-methionine (SAM) and associated metabolites at toxic ASNase doses prompted the hypothesis that SAM boosting as a nutritional supplement may mitigate ASNase toxicity (**Figure 3E**).

In the lipidomic data, notable trends included up-regulation of gangliosides (GM3) (**Figure 3F**) and inflammatory lipids such as ceramide (Cer), TG, and DG at toxic doses of ASNase (**Figure 3G**). Cer have been well-documented to promote inflammation through activation of diverse signaling pathways^18^. The accumulation of TGs and DGs has been shown to trigger inflammatory responses in pancreatic tissue^19^. These changes coincided with suppression of polyunsaturated fatty acids (PUFAs) at toxic doses of ASNase (**Figure 3H**). These observations prompt the hypothesis that ASNase-induced pancreatitis, which occurs in up to 18% of patients and can be a deadly side effect, may be mitigated through supplementation with PUFAs^20^.

Finally, KEGG pathway analysis of proteins differentially modulated by toxic doses of ASNase revealed up-regulation of cholesterol metabolism, cholesterol metabolic process, cholesterol homeostasis, cholesterol efflux, and reverse cholesterol transport proteins (**Figure 3I**), consistent with hypertriglyceridemia as a known toxicity of ASNase. Toxic doses of ASNase also caused up-regulation of proteins associated with hemostasis, blood coagulation, and clotting cascade with concomitant downregulation of complement activation and complement and coagulation cascades, all of which are consistent with venous thromboembolism (VTE) as a side effect of ASNase. Toxic ASNase doses also upregulated “neutrophil degranulation, which aligns with the up-regulation of inflammatory lipids noted previously.

We were particularly interested in the up-regulation of cell proliferation proteins at the sub-toxic but not toxic doses, because this observation suggests that insufficient concentration of ASNase can accelerate cancer proliferation, which we have observed with our use of a glutaminase-free Q59L mutant of ASNase^21^. Finally, down-regulation of humoral immune response proteins align with the known immunosuppressive side effects of ASNase.

Next, integrating metabolomic, lipidomic, and proteomic data using a systems biology graphical notation (SBGN) revealed that ASNase treatment orchestrates complex molecular networks that differ markedly between sub-toxic and toxic doses. The integrated molecular network analysis (**Figure 4A**) demonstrates extensive cross-talk between metabolic pathways, lipid metabolism, and protein responses. At sub-toxic doses (2,500 IU/kg), cells maintain homeostasis through adaptive responses, shown by coordinated regulation of metabolites (amino acids and energy metabolism), phospholipids (PC, PS, PE, DG, TG, and ceramide), and protective proteins. However, at toxic doses (10,000 and 20,000 IU/kg), this balance is disrupted through multiple interconnected pathways, as evidenced by the inflammatory lipid network signatures (**Figure 4B**). Key inflammatory mediators, including ADIPOQ, APCS, and CDH5, exhibited progressive upregulation with increasing ASNase dose.

**Figure 4.**
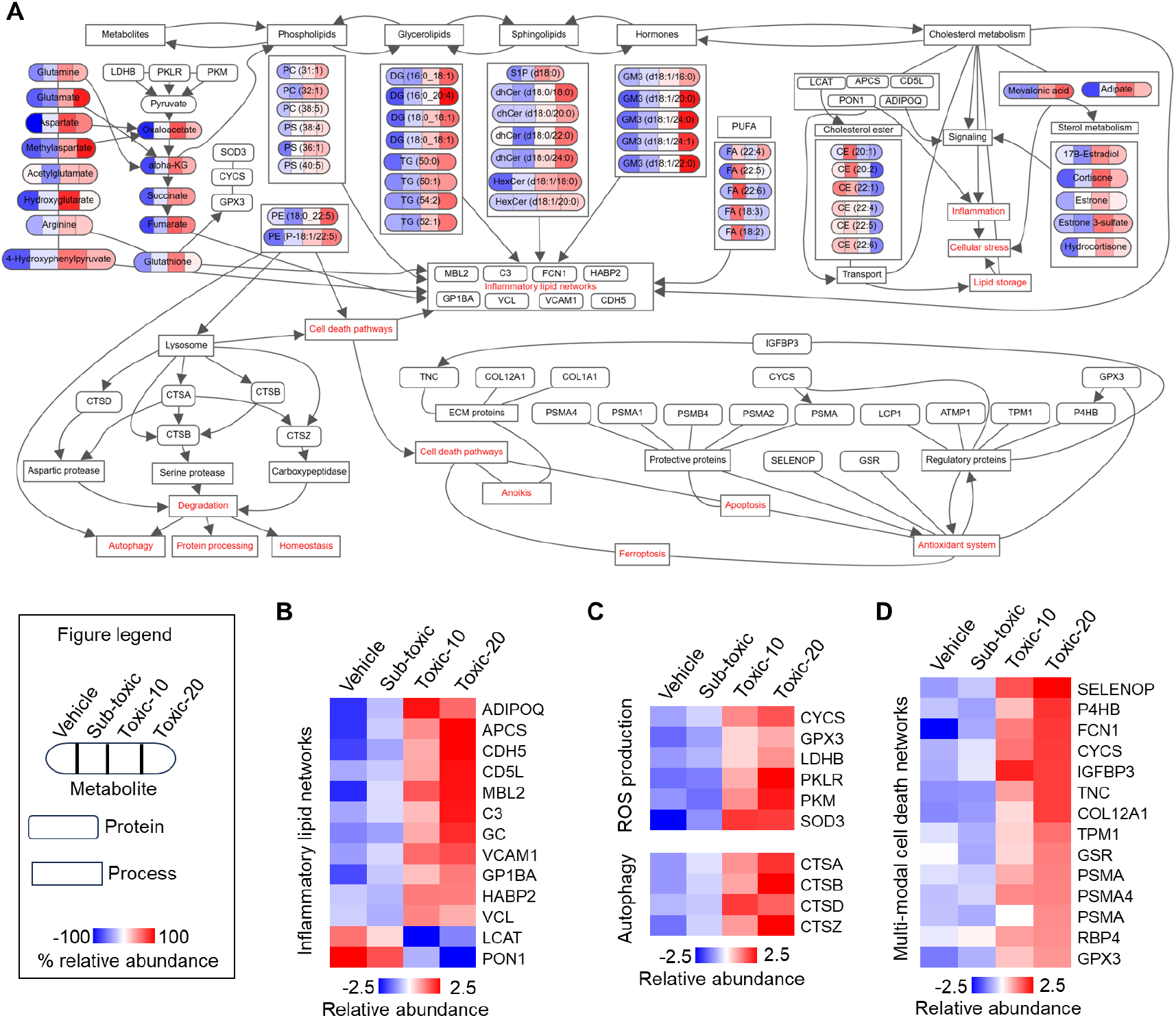
Integrated molecular networks showing differential effects of sub-toxic and toxic ASNase dosages. (**A**) Integrated molecular network visualization showing known connections between metabolites, lipids, and proteins affected by ASNase treatment. (B) Heatmap showing differential modulation of inflammatory lipid networks across treatment groups, highlighting key proteins involved in lipid metabolism and inflammation (ADIPOQ, APCS, CDH5, etc.). (C) Heatmap depicting changes in oxidative phosphorylation (OXPHOS), reactive oxygen species (ROS) production, and autophagy-related proteins, showing progressive activation with increasing ASNase doses. (D) Heatmap illustrating multi-modal cell death protein signatures, including proteins involved in apoptosis, oxidative stress response (SELENOP, GPX3), and cell death regulation (FCN1, CYCS), demonstrating dose-dependent activation of cell death pathways.

Proteins involved in ROS production (CYC and GPX3), OXPHOS (LDHB, PKLR, and PKM), and autophagy (CTSA, CTSB, CTSD, and CTSZ) were significantly elevated at toxic ASNase doses (**Figure 4C**). Finally, we observed evidence of activation of multi-modal cell death pathways (**Figure 4D**), marked by upregulation of death-associated proteins (SELENOP, FCN1, and CYCS) and stress response factors (TPM1 and GSR). Overall, the observed molecular changes were consistent with known clinical ASNase toxicities, including pancreatitis, thrombosis, hyperglycemia, and immunosuppression (**Figure 4**).

## CONCLUSION

We present a streamlined LC-MS/MS method for quantitative analysis of 500 metabolites, 750 lipids, and 375 proteins from a single 20 µL plasma sample. The remarkable precision achieved presents a viable alternative or complement to non-targeted workflows aimed at elucidating drug mechanisms and biomarkers of drug sensitivity or resistance.

Our application of the method unveiled new insights into the mechanisms of ASNase toxicity. At toxic doses, ASNase compromised antioxidant systems and activated inflammatory networks and ganglioside production. Those observations prompt future testing of therapeutic interventions including antioxidant supplementation, PUFA supplementation, or anti-inflammatory agents to potentially increase the therapeutic index of ASNase (**Figure 5**). Future studies should also focus on developing biomarker-guided, therapeutic drug monitoring protocols.

**Figure 5.**
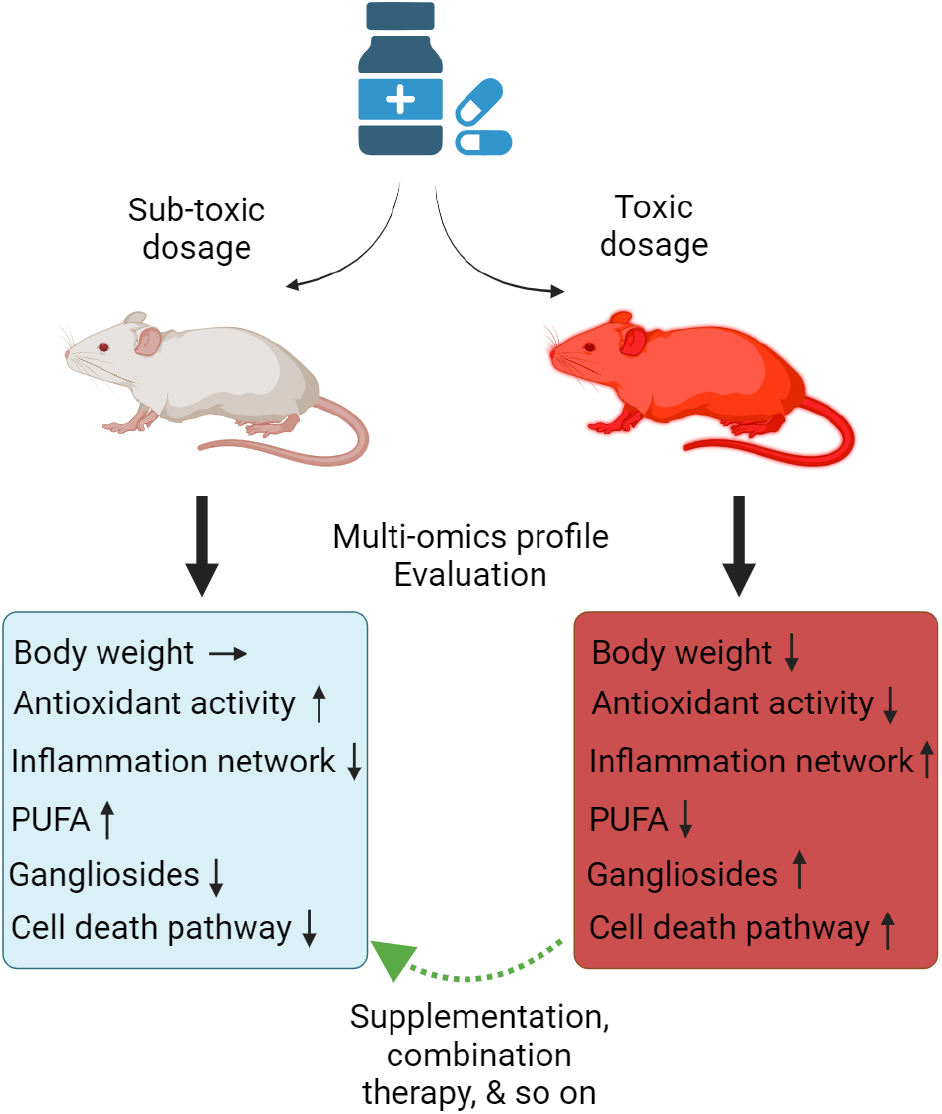
Potential strategies for mitigating dysregulation of the multi-omic response to toxic ASNase dosages.

In summary, we demonstrate the potential of the workflow to produce new insights into drug mechanisms and targetable biomarkers and pathways for decreasing drug toxicity. It can be applied to a range of multi-omic studies in mice as well as other basic and translational systems beyond our particular application to the toxicity of ASNase.

## ASSOCIATED CONTENT

Supporting Information

## AUTHOR INFORMATION

### Corresponding Authors

Philip L. Lorenzi, The University of Texas MD Anderson Cancer Center, 7435 Fannin Street, Room 2SCR3.3029, Unit 0950, Houston, TX 77054.

Phone: 713-792-9999.

Iqbal Mahmud, The University of Texas MD Anderson Cancer Center, 7435 Fannin Street, Room 2SCR3.3029, Unit 0950, Houston, TX 77054.

Phone: 713-792-9999.

## Author Contributions

IM, WC, KEY, GCvdB, JS, & PLL – Study design. IM - writing the original draft. KEY, GCvdB, & WC contributed to writing. KEY, GCvdB, and WC – sample preparation. IM, CS, DC, YL, SBM, LW – data collection and analysis. IM, KEY, & PLL – data interpretation and visualization. IM, WC, KEY, GCvdB, JNW, JS, & PLL – reviewed and edited the manuscript.

## Notes

IM, WC, DC, YL, JNW, & PLL declare no competing financial interest. KEY, GCvdB, CS, LW, SBM, and JS are employed by Agilent Technologies.

## ACKNOWLEDGMENT

The Metabolomics Core Facility is supported by NIH grants S10OD012304-01 and P30CA016672

## Supplementary figures

**Figure S1.**
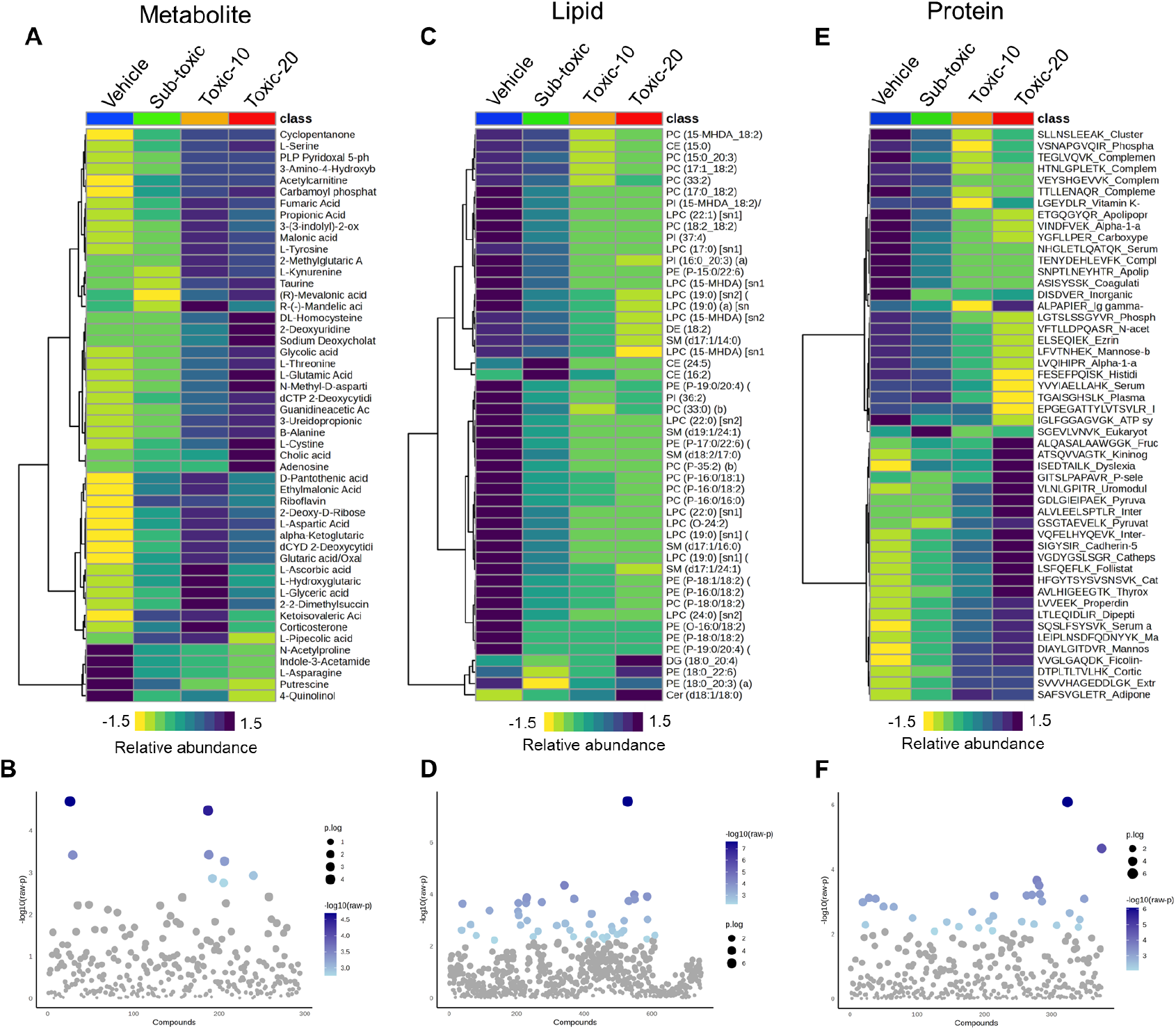
Relative abundance of biomarker at multi-omics level. Relative abundances are shown for top 50 metabolite (A), lipid (C), and protein (E). One-way ANOVAs are shown for metabolite (B), lipid (D), and protein (F).

**Figure S2.**
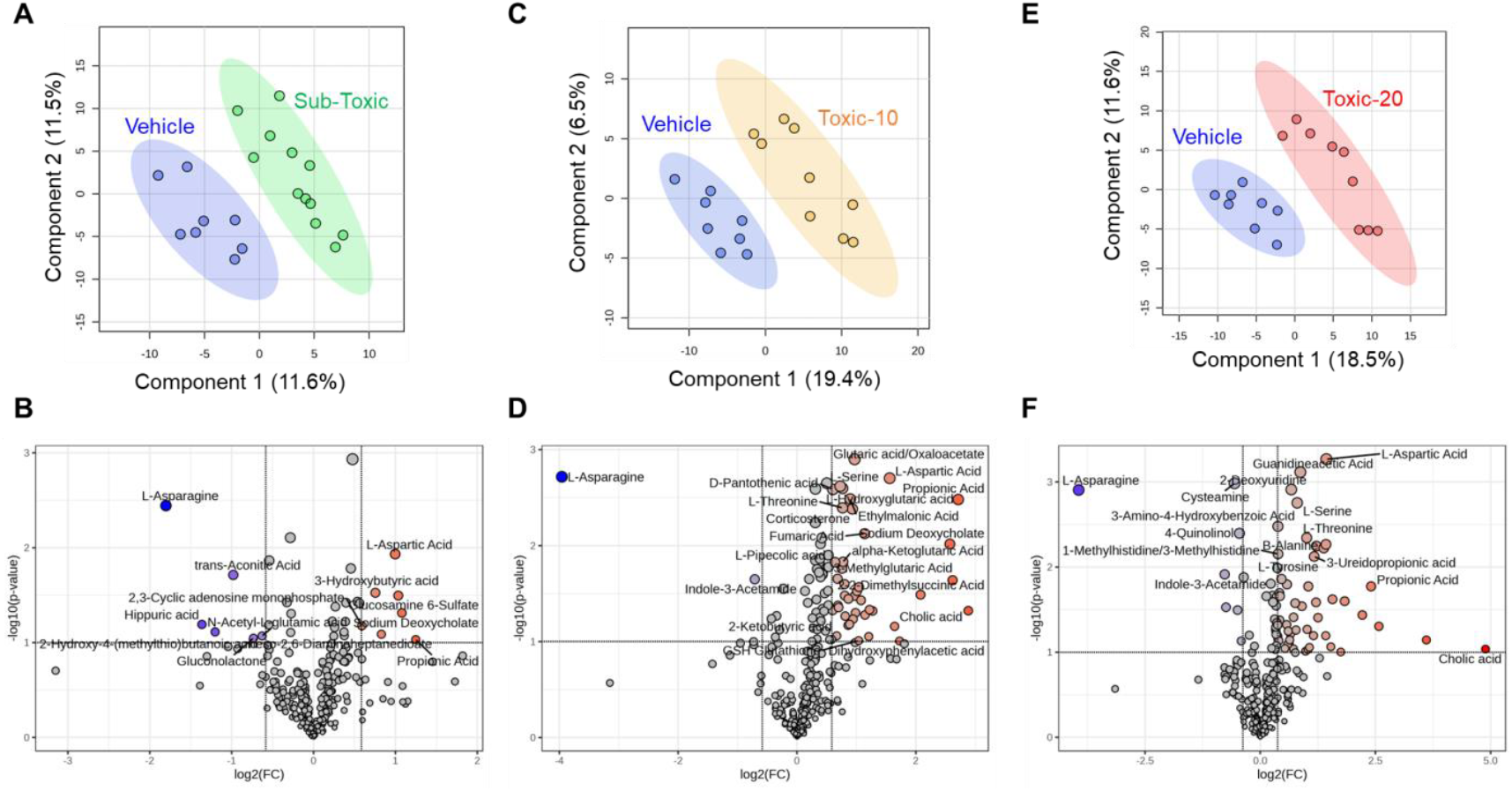
ASNase modulates global metabolite profile. Pair-wise metabolic profile is shown for vehicle versus sub-toxic (A), vehicle versus Toxic-10 (C), and vehicle versus Toxic-20 (E) dosages. Significantly modulated metabolite profile is shown for sub-toxic (B), Toxic-10 (D), and Toxic-20 (F) compared to vehicle.

**Figure S3.**
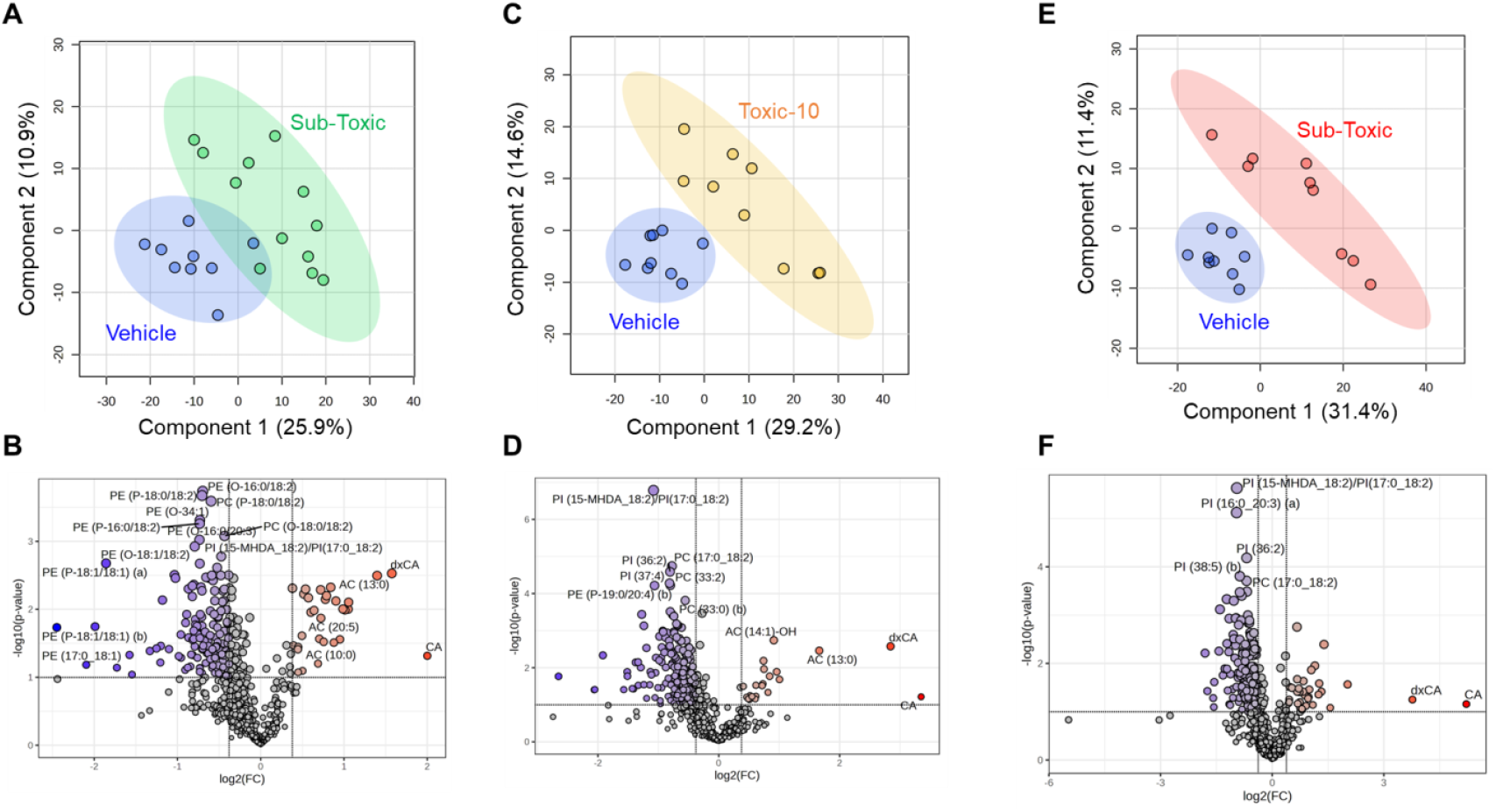
ASNase modulates global lipid profile. Pair-wise lipid profile is shown for vehicle versus sub-toxic (A), vehicle versus Toxic-10 (C), and vehicle versus Toxic-20 (E) dosages. Significantly modulated lipid profile is shown for sub-toxic (B), Toxic-10 (D), and Toxic-20 (F) compared to vehicle.

**Figure S4.**
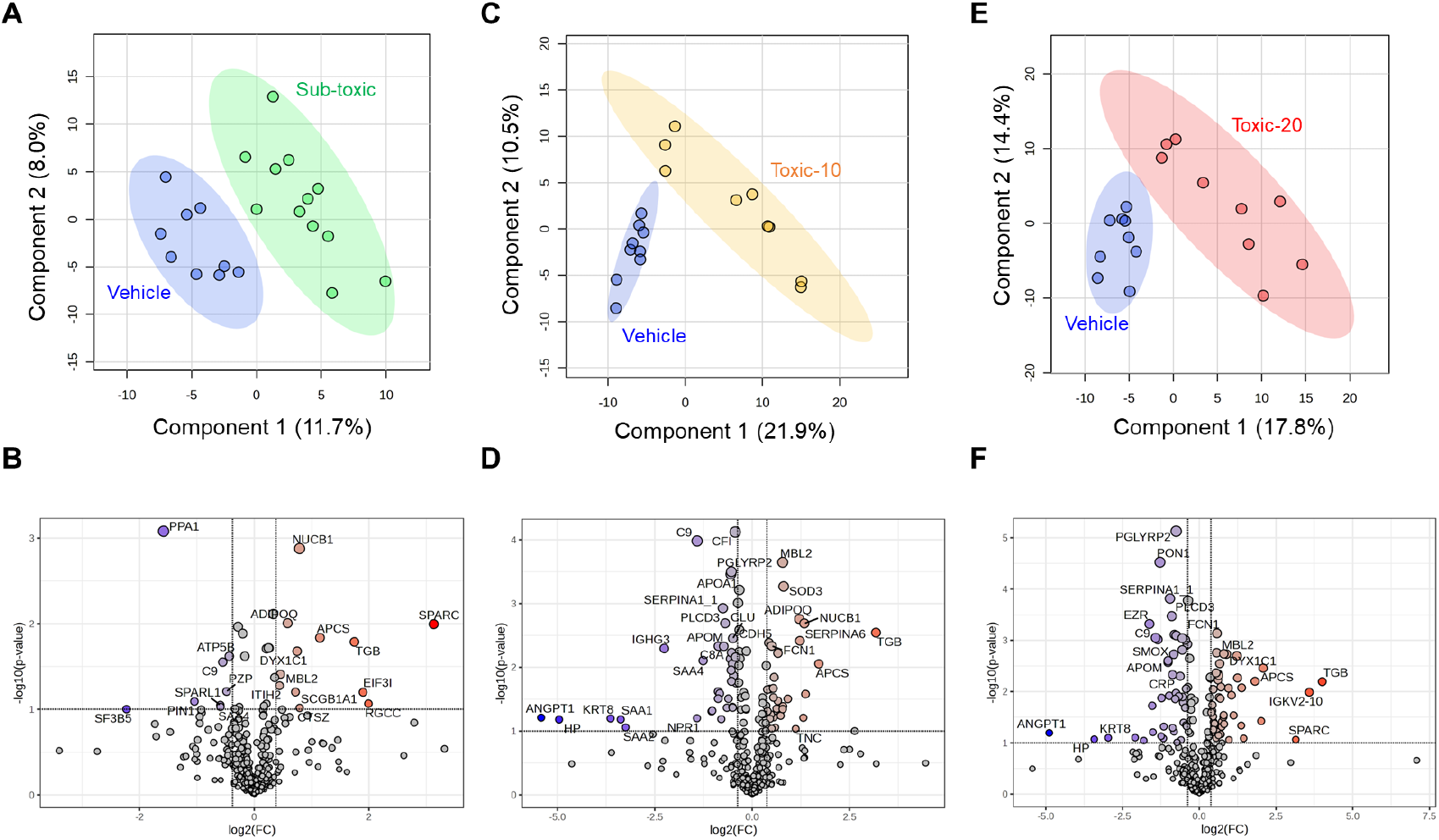
ASNase modulates global protein profile. Pair-wise protein profile is shown for vehicle versus sub-toxic (A), vehicle versus Toxic-10 (C), and vehicle versus Toxic-20 (E) dosages. Significantly modulated protein profile is shown for sub-toxic (B), Toxic-10 (D), and Toxic-20 (F) compared to vehicle.

